# DDX41 recognizes RNA/DNA retroviral reverse transcripts and is critical for *in vivo* control of MLV infection

**DOI:** 10.1101/312777

**Authors:** Spyridon Stavrou, Alexya Aguilera, Kristin Blouch, Susan R. Ross

## Abstract

Host recognition of viral nucleic acids generated during infection leads to the activation of innate immune responses essential for early control of virus. Retrovirus reverse transcription creates numerous potential ligands for cytosolic host sensors that recognize foreign nucleic acids, including single-stranded RNA (ssRNA), RNA/DNA hybrids and double stranded DNA (dsDNA). We and others recently showed that the sensors cyclic GMP-AMP synthase (cGAS), dead-box helicase 41 (DDX41) and members of the Aim2-like receptor (ALR) family participate in the recognition of retroviral reverse transcripts. However, why multiple sensors might be required and their relative importance in *in vivo* control of retroviral infection is not known. Here we show that DDX41 primarily senses the DNA/RNA hybrid generated at the first step of reverse transcription, while cGAS recognizes dsDNA generated at the next step. We also show that both DDX41 and cGAS are needed for the anti-retroviral innate immune response to MLV and HIV in primary mouse macrophages and dendritic cells (DC). Using mice with macrophage- or -specific knockout of the DDX41 gene, we show that DDX41 sensing in DCs but not macrophages was critical for controlling *in vivo* MLV infection. This suggests that DCs are essential *in vivo* targets for infection, as well as for initiating the antiviral response. Our work demonstrates that the innate immune response to retrovirus infection depends on multiple host nucleic acid sensors that recognize different reverse transcription intermediates.

**Importance:** Viruses are detected by many different host sensors of nucleic acid, which in turn trigger innate immune responses, such as type I IFN production, required to control infection. We show here that at least two sensors are needed to initiate a highly effective innate immune response to retroviruses – DDX41, which preferentially senses the RNA/DNA hybrid generated at the first step of retrovirus replication and cGAS, which recognizes double-stranded DNA generated at the 2^nd^ step. Importantly, we demonstrate using mice lacking DDX41 or cGAS, that both sensors are needed for the full antiviral response needed to control *in vivo* MLV infection. These findings underscore the need for multiple host factors to counteract retroviral infection.

## Introduction

Retroviruses are major causes of disease in animals and humans. The initial immune response to retroviruses is critical to the ability of organisms to clear infection, because once viral DNA integrates into the host chromosomes, persistent infections arise, leading to immunodeficiencies, cancers and other pathologies. The genomes of mammals and other species encode many genes that restrict infectious retroviruses. Among the host anti-retroviral factors, APOBEC3 proteins play a major role in restricting retrovirus infection, by cytidine deamination of retroviral DNA and by blocking early reverse transcription (1-7).

The retrovirus RNA genome is converted by the viral reverse transcriptase (RT) enzyme first to RNA/DNA hybrids using a tRNA to prime DNA synthesis and then to dsDNA. Reverse transcription thus creates potential ligands for host sensors that recognize foreign nucleic acids. Cellular recognition of these retroviral reverse transcripts activates the innate immune response. For example, depletion of the host cytosolic DNA exonuclease Three Prime Repair Exonuclease 1 (TREX1), a DNA exonuclease, increases the type I interferon (IFN) response to HIV and MLV infection (4, 8, 9). The TREX1-sensitive retroviral reverse transcripts are recognized by cellular DNA sensors such as cGAS, DDX41 and ALR family members such as IFN-Induced 16 (IFI16) in humans and IFI203 in mice (9-13).

cGAS produces the second messenger cyclic GMP-AMP (cGAMP) upon DNA binding, which binds and activates Stimulator of IFN Genes (STING) (14-16). STING then translocates from the endoplasmic reticulum to a perinuclear compartment and activates TANK-binding kinase 1 (TBK1) which phosphorylates the transcription factor IFN regulatory factor 3 (IRF3) which in turn enters the nucleus where it induces type 1 IFN transcription (17-19). DNA binding to DDX41 and the ALRs also induces type I IFN production via the STING pathway (20). It is not understood how DDX41, which belongs to a family of DEAD box helicase-containing genes commonly thought to bind RNA, participates in the recognition of nucleic acid. Familial and sporadic mutations in human DDX41 lead to acute myeloblastic leukemia and myelodysplastic syndromes, suggesting that it also functions as a tumor suppressor (21, 22).

While many studies have shown that the loss of any one of these factors decreases the STING-mediated IFN response to cytosolic DNA, it is not known why there are multiple sensors that converge on the same pathway, particularly *in vivo*. Here we show that DDX41 recognizes the RNA/DNA intermediate generated by reverse transcription and that DDX41 and cGAS act additively to increase the IFN response and limit retroviral infection *in vivo*. Moreover, using mice with cell type-specific knockout of DDX41, we show that DCs and not myeloid-derived cells are likely the major sentinel cell targets of *in vivo* infection. These studies reveal why multiple nucleic acid sensors are needed to control retroviral infection and underscore the importance of studying their role in *in vivo* infection.

## Results

### DDX41, IFI203 and cGAS play independent but additive roles in the response to MLV infection

We showed previously that MLV infection caused a rapid increase in IFNβ RNA levels in murine macrophages that is sensitive to the RT inhibitor ziduvodine and that Trex1 knockdown further increased this response (4, 9). We also showed that depletion of DDX41, IFI203 or cGAS diminished the IFNβ response with and without Trex1 knockdown, that all three molecules bound MLV reverse transcribed DNA and that IFI203 and DDX41 bound to each other and STING, but not to cGAS (9). These data suggested that IFI203 and DDX41 work together in a complex to sense reverse transcripts.

We hypothesized that DDX41/IFI203 and cGAS play additive but non-redundant roles in the STING/IFNβ activation pathway. To determine if DDX41/IFI203 and cGAS acted synergistically to generate an anti-MLV response, we tested the effects of DDX41, IFI203, and STING knockdown in bone marrow-derived macrophages (BMDMs) and DCs (BMDCs) isolated from cGAS knockout (KO) mice that also lacked APOBEC3; APOBEC3 depletion leads to increased reverse transcript levels and higher levels of IFN induction and thus greater assay sensitivity (4, 9). After siRNA-mediated knockdown, the cells were infected with MLV and the IFN response determined at 2 hpi, the time of maximum response (4, 9). Despite the lack of cGAS in these cells, MLV infection induced higher levels of IFNβ RNA that was further increased by Trex1 knockdown (compare mock to control to Trex1 siRNA in Fig. 1A), suggesting that additional sensors of retroviral nucleic acid exist in sentinel cells. Ddx41 or Ifi203 knockdown in cGAS KO BMDMs and BMDCs diminished the type I IFN response to the same level as Sting knockdown (Fig. 1A). These data show that the full STING-dependent type I IFN response to MLV reverse transcripts requires both DDX41/IFI203 and cGAS.

**Fig. 1.**
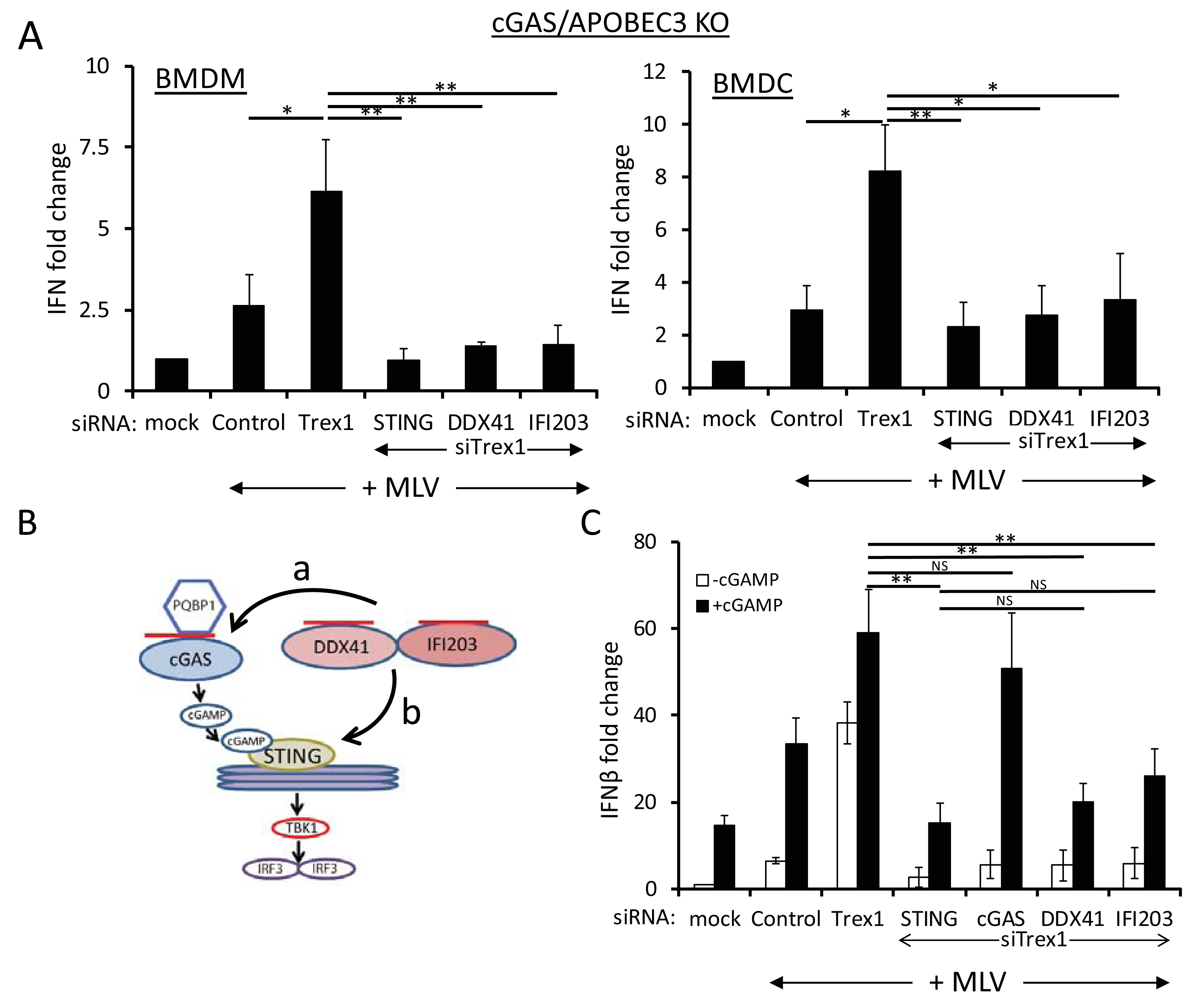
DDX41 and IFI203 work together with cGAS for the maximal anti-viral response. A) Knockdown of STING, DDX41 and IFI203 in cGAS/APOBEC3 double knockout BMDMs and BMDCs. Cells were transfected with the indicated siRNAs and 48hrs later cells were infected with MLV. At 2 hpi, the cells were harvested and examined for IFNβ RNA levels. Knockdown verification of the genes is shown in Fig. S1A. Values are shown as mean ± STDs of three experiments, each with macrophages and DCs from a different mouse. P values were determined by unpaired T-tests (NS, not significant; *, p≤ 0.05; **, p≤ 0.01). B) Diagram shows the cGAS-cGAMP-STING pathway. The dotted line represents the possible points of DDX41 action; cGAMP addition would rescue DDX41 knockdown if it acted at (a) but not if DDX41 acted at (b) in the pathway. The red lines represent viral reverse transcripts. C) cGAMP rescues cGAS but not STING, DDX41 or IFI203 knockdown. NR9456 macrophages were transfected with the indicated siRNAs and 24 hr later, transfected with cGAMP. At 18 hr post-cGAMP treatment, the cells were infected with MLV; IFNβ RNA levels measured at 2 hpi. Values are shown as mean ± STDs of three experiments. Knockdown verification of the genes is shown in Fig. S1B. Mock denotes mock-infected cells.

The factor PQBP1 binds retroviral DNA upon infection and functions upstream of cGAS, since cGAMP addition to PQBP1-or cGAS-depleted cells restores the type I IFN response (23) (diagram in Fig. 1B). To determine if DDX41 worked upstream of cGAS, we tested whether cGAMP also would rescue the IFN response in Ddx41/Ifi203 knockdown cells. siRNA-mediated depletion of Ifi203 and Ddx41 plus Trex1 was carried out in NR9456 mouse macrophage cells; cGAS and STING depletion served as positive and negative controls, respectively. At 24 hr post-siRNA transfection, the cells were left untreated or transfected with cGAMP for 18 hr and then infected with MLV for 2 hr. cGAMP addition did not restore the IFNβ response in DDX41-, IFI203- or STING-depleted cells (Fig. 1C). As expected, addition of cGAMP restored the IFNβ response in cGAS-depleted cells (Fig. 1C). Taken together with our previously published results, these data suggested that DDX41/IFI203 functions independently of cGAS to activate the STING pathway (pathway b in Fig. 1B) and that the induction of IFN by the two sensors is additive.

### DDX41 is a cytosolic sensor that acts upstream of IRF3 and TBK1

DDX41 is found in both the nucleus and cytoplasm (9). The ALR IFI16 senses herpes simplex virus DNA in the nucleus and then migrates to the cytoplasm where it signals through STING (24, 25). The ultimate product of reverse transcription is a dsDNA that is transported into the nucleus and integrates into the chromosomes. Unintegrated retroviral DNA persists in the nucleus as 1- or 2-LTR circles. To determine if DDX41 sensed nuclear retroviral dsDNA, we treated cells with the integrase inhibitor raltegravir, which increases nuclear unintegrated viral dsDNA levels, and examined the IFN response after MLV infection. Although raltegravir treatment dramatically increased the levels of unintegrated nuclear viral DNA, evidenced by abundant 2-LTR circle formation, this treatment had no effect on IFNβ induction (Fig. S2A), supporting DDX41 sensing of retroviral reverse transcription products predominantly in the cytoplasm.

To test whether DDX41 functioned downstream of STING, TBK1 or IRF3 (Fig. 1B), we siRNA-depleted DDX41, cGAS or STING in NR9456 macrophages, infected them with MLV and examined IRF3 (Ser396) and TBK1 (Ser172) phosphorylation at 2 hr post-infection (hpi); lipopolysaccharide (LPS) treatment served as a positive control. While phospho-TBK1 and -IRF3 were induced in LPS-treated BMDMs and in Trex1-depleted BMDMs in response to MLV, siRNA depletion of Ddx41, cGas or Sting ablated virus-induced TBK1 and IRF3 phosphorylation (Fig. 2). Taken together, these data suggest that DDX41 works in the cytoplasm upstream of STING to induce IFN.

**Fig. 2.**
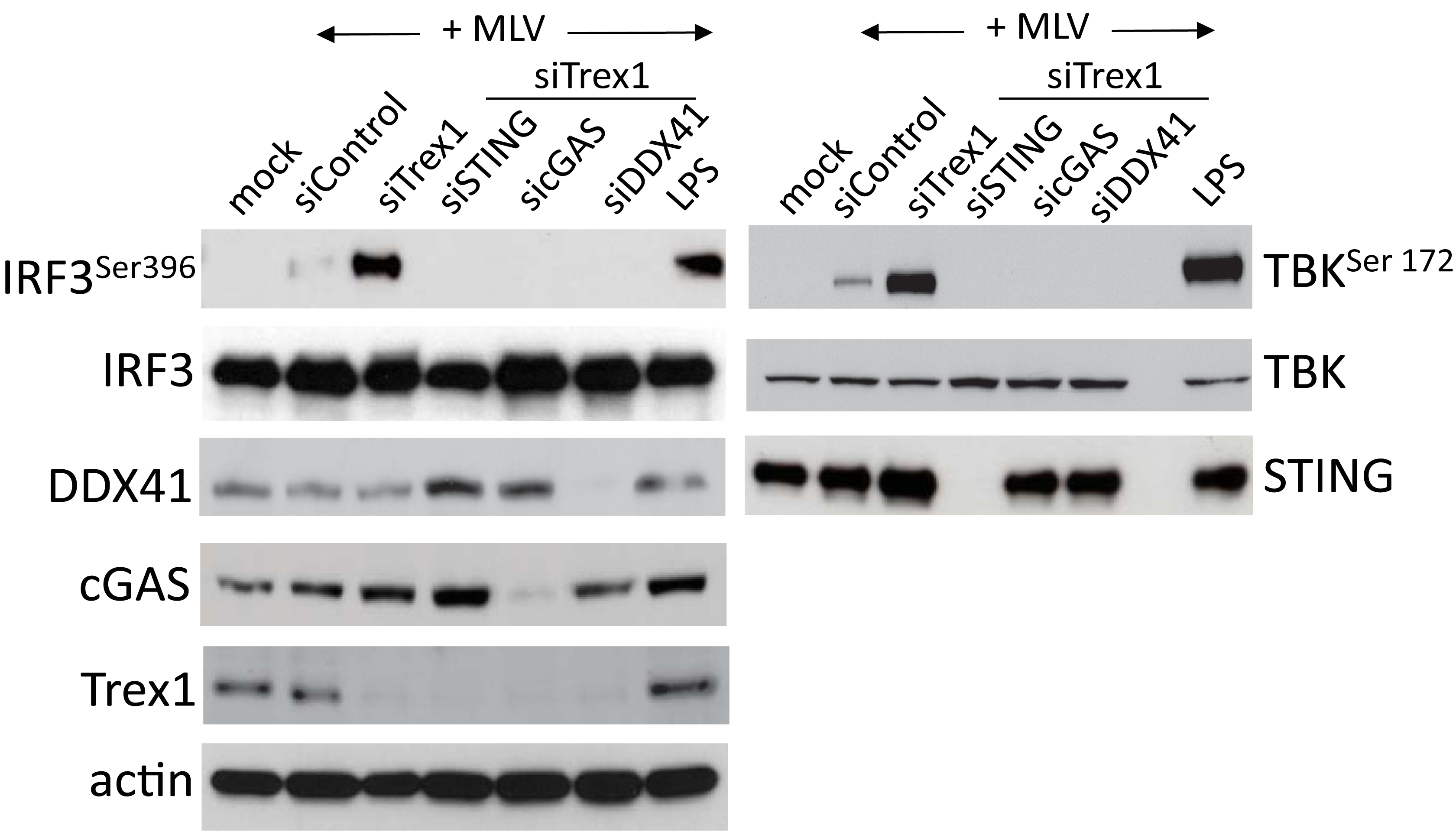
DDX41 acts upstream of IRF3 and TBK1. IRF3 (left panel) and TBK (right panel) phosphorylation induced by MLV infection requires DDX41, cGAS and STING. NR9456 cells were transfected with the indicated siRNAs as well as Trex1 siRNA and 48hr later, infected with MLV for 2 hr. Control cells were infected but received only control siRNA. The LPS treatments were for 6 hr. Equal amounts of protein from the cells were analyzed using the indicated antibodies. Mock denotes mock-infected cells. The TBK1 and IRF3 experiments were performed twice. Shown are representative western blots.

### DDX41 recognizes RNA/DNA hybrid reverse transcription intermediates

Retroviruses generate several replication intermediates which could be sensed as foreign – tRNA-bound DNA/RNA hybrids, ssDNA and dsDNA. We used three approaches to examine which reverse transcription products were sensed by DDX41. First, to determine if DDX41 or cGAS bound to tRNA primer-containing reverse transcription intermediates, 293T cells stably expressing the MLV receptor MCAT1 were transiently transfected with DDX41 or cGAS constructs, infected with MLV and pull-down experiments were performed. After pull-down, DNA was isolated from half of each sample and subjected to PCR amplification with primers that detect early reverse transcripts (strong-stop primers P_R_ and P_U5_), while cDNA was prepared from the remaining half and amplified with P_R_ and a 3’ primer specific to tRNA^pro^ (P_tRNA_), the tRNA used by MLV RT to prime reverse transcription (Fig. 3A). DDX41 bound to >2-fold more tRNA^pro^-containing reverse transcripts, while DDX41 and cGAS equally precipitated a product that amplified strong stop DNA (Fig. 3B).

**Fig. 3.**
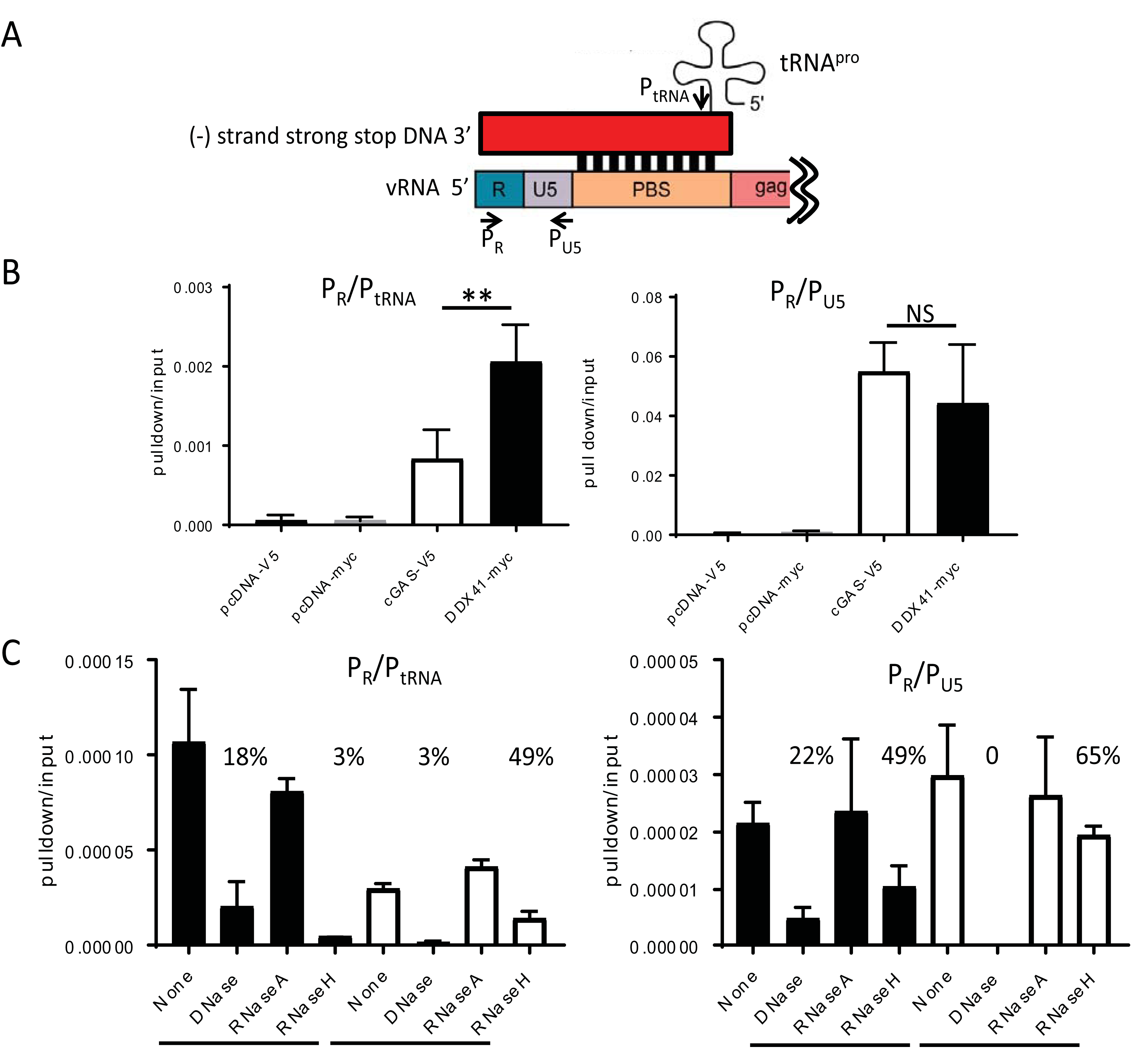
DDX41 preferentially binds RNA/DNA hybrids. A) Diagram of the tRNA/LTR (P_R_ and P_tRNA_) and strong stop primers (P_R_ and P_U5_) used to amplify the bound nucleic acid in (B). Red box represents the newly synthesized viral DNA; shown below is the viral RNA. Abbreviations: vRNA, viral RNA; PBS, primer binding site. B) DNA pulldown assays with extracts from 293T MCAT-1 cells transfected with the indicated constructs and infected with MLV. Shown is the mean of 4 independent experiments ± STD. **, p≤ 0.01 (unpaired T-test); NS, not significant. C) DNA pulldown assays were conducted as in (B), except that prior to the reverse transcription/RT-qPCR, the nucleic acids were treated with the indicated nucleases. Shown is the average of two experiments done in triplicate. The numbers above the columns show percent nucleic acid pulldown relative to no nuclease.

Second, we treated the DDX41- and cGAS-bound nucleic acids with RNaseH, which degrades RNA in DNA/RNA hybrids as well as the tRNA primer, DNaseI, which cleaves dsDNA 100- and 500-fold better than RNA/DNA hybrids and ssDNA, respectively, and RNaseA, which degrades ssRNA under high salt conditions. DDX41 again more efficiently precipitated the RNA/DNA hybrid and RNaseH treatment reduced the amount of DDX41-precipitated nucleic acid to 3%. In contrast, RNaseH digestion only modestly affected cGAS-pulldown of the product amplified with the P_R_/P_tRNA_ primer pair, suggesting that DDX41 preferentially bound the RNA/DNA hybrid while cGAS bound to tRNA primer-bound dsDNA generated after strand translocation (Fig. 3C, left panel; see diagram in Fig. 4A). In support of this, DNaseI treatment abolished cGAS-mediated pulldown of both the P_R_/P_tRNA_- and P_R_/P_U5_-amplifiable products, while DDX41-mediated precipitation of nucleic acid (Fig. 3C, left panel and right panels) was affected to a lesser extent. RNaseA digestion in high salt had no effect on any of the pulldowns.

**Fig. 4.**
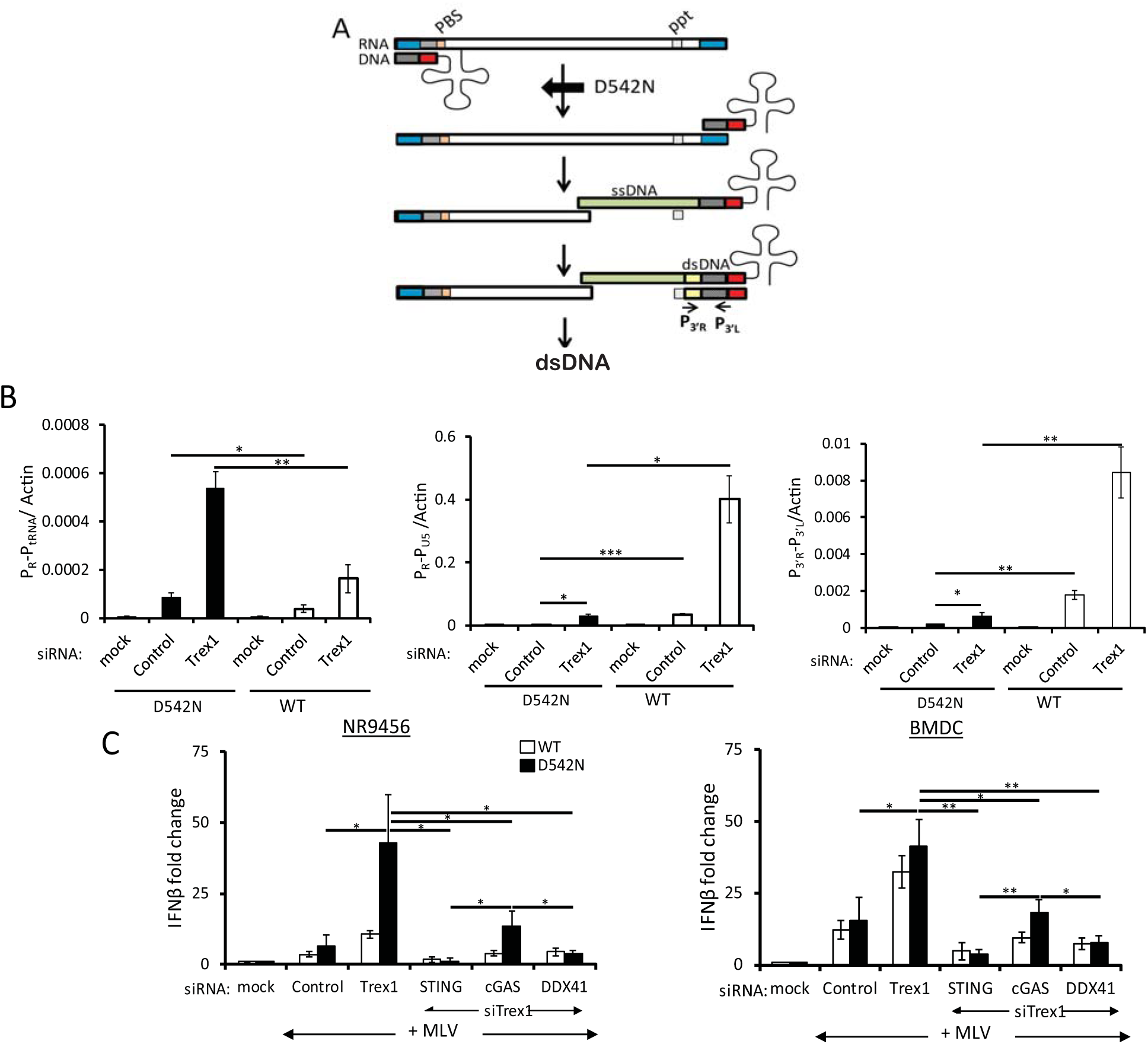
MLV^D542N^ reverse transcripts are sensed by DDX41. A) Diagram of the early stages of reverse transcription. The RNAseH^D542N^ mutation allows initial reverse transcription but because of the loss of RNAseH activity, strong stop DNA cannot translocate to the 3’ end of the viral RNA to initiate transcription of the full length viral dsDNA; the tRNA primer is also not degraded. P_3’R_ and P_3’L_ denote the 3’LTR primers used in panel B. Abbreviations: PBS, primer binding site; ppt, polypurine tract; ssDNA, single-strand DNA; ds, double-strand DNA. B) NR9456 cells treated with the indicated siRNAs were infected with D542N or wildtype virus and 2 hpi, RNA was subjected to reverse transcribed-qPCR with primers to the tRNA-containing RNA/DNA hybrid (P_R_ and P_tRNA,_ Fig. 3A), while DNA was subjected to qPCR with strong-stop DNA (P_R_ and P_U5_,Fig. 3A) or late reverse transcripts (P_3’R_ and P_3’L_ primers, Fig. 4A). Shown is the mean ± STDs of 3 independent experiments. *, p≤ 0.05; **, p≤ 0.01; ***, p≤ 0.001 (unpaired T-test). C) Recognition of RNA-DNA hybrids requires DDX41. NR9456 cells (left panel) or BMDCs (right panel) were transfected with the indicated siRNAs, infected with wild type MLV or MLV^D542N^ for 2 hr and the levels of IFNβ RNA were measured. Shown is the mean ± STD of 3 independent experiments. *, p≤ 0.05; **, p≤ 0.01; ***, p≤ 0.001 (unpaired T-test). Knockdown of the genes is shown in Fig. S3. Mock denotes mock-infected cells.

Finally, we used a viral mutant lacking RNaseH activity. During reverse transcription, RT’s RNaseH moiety degrades the positive strand RNA genome after the synthesis of minus strand DNA (Fig. 4A) (26). RNaseH mutations attenuates the RNaseH function without diminishing the polymerase activity. RNaseH^D542N^ synthesizes tRNA^pro^-primed (–) strand strong stop DNA while retaining ^~^10% the wild type levels of RNaseH enzymatic activity. As a result, the RNA remains “frozen” in a DNA/RNA hybrid and (-) strand strong stop DNA does not efficiently translocate to the 3’ end of the viral RNA to initiate full-length (-) strand DNA synthesis (Fig. 4A) (27, 28). We engineered the D542N mutation into a MLV molecular clone (MLV^D542N^) and used this virus to infect NR9456 cells. MLV^D542N^ generated almost 3-fold more reverse transcription products retaining the tRNA primer than did the wild type virus, reflecting its poorer ability to translocate the negative strand strong-stop DNA to the 5’ end of the RNA and degrade the tRNA primer, and its known increased DNA polymerase activity relative to wild type virus (P_R_-P_tRNA_, Fig. 4B) (27, 29). The mutation dramatically attenuated reverse transcription detected with the strong-stop (P_R_-P_U5_) primers compared to wild type virus, since these primers detect R-U5 DNA present in (-) and (+) strand strong stop as well as full-length (-) strand DNA; late reverse transcription (P_3’R_-P_3’L_) products were also reduced compared to wild type virus (Fig. 4B). Interestingly, Trex1 depletion led to increases in reverse transcription products retaining tRNA from both the wild type and MLV^D542N^ viruses, suggesting that negative-strand strong-stop DNA is also degraded by this cellular exonuclease (Fig. 4B); it has been previously shown that TREX1 degrades ssDNA and dsDNA and DNA in RNA/DNA hybrids (8, 30, 31).

To determine whether DDX41 or cGAS was better able to recognize the early DNA/RNA reverse transcription product, we infected NR9456 cells or primary BMDCs with MLV^D542N^ after treatment with Trex1 siRNA alone, or in combination with Sting, Ddx41 or cGas siRNAs. MLV^D542N^ caused about a 5- and 2-fold increase in the IFN response compared to wild type virus in the Trex1 siRNA-treated and untreated NR9456 cells, respectively (left panel, Fig. 4C). DDX41 knockdown diminished the response to both viruses to the same levels as seen with STING knockdown in both NR9456 cells and primary BMDCs (Fig. 4C). Depletion of cGAS reduced but did not completely abrogate the TREX1-dependent response to MLV^D542N^ in NR9456 or primary BMDCs (Fig. 4C). The response to the RNaseH mutant virus in cGAS-deficient cells was likely due to DDX41-mediated recognition of the RNA/DNA hybrid in these cells (Fig. 4C).

The results from these 3 complementary approaches indicate that DDX41 preferentially senses the RNA/DNA hybrid generated during the earliest stage of reverse transcription while cGAS preferentially recognizes dsDNA generated at the next step.

### DDX41 is required for the IFN response in both macrophages and DCs

Macrophages and DCs have both been implicated in the anti-retroviral innate immune response. We examined DDX41 expression in BMDMs and BMDCs and found that it was expressed in both cell types at both the RNA and protein level (Fig. 5A). Interestingly, in contrast to DDX41 expression, cGas and Ifi203 RNA levels and cGAS protein levels were significantly higher in wild type BMDMs than in BMDCs, suggesting that the sensors used to detect nucleic acid might be cell type-specific (Fig. 5A). The lack of anti-IFI203-specific antisera prevented us from determining whether its protein levels in BMDMs and BMDCs reflected the RNA levels.

**Fig. 5.**
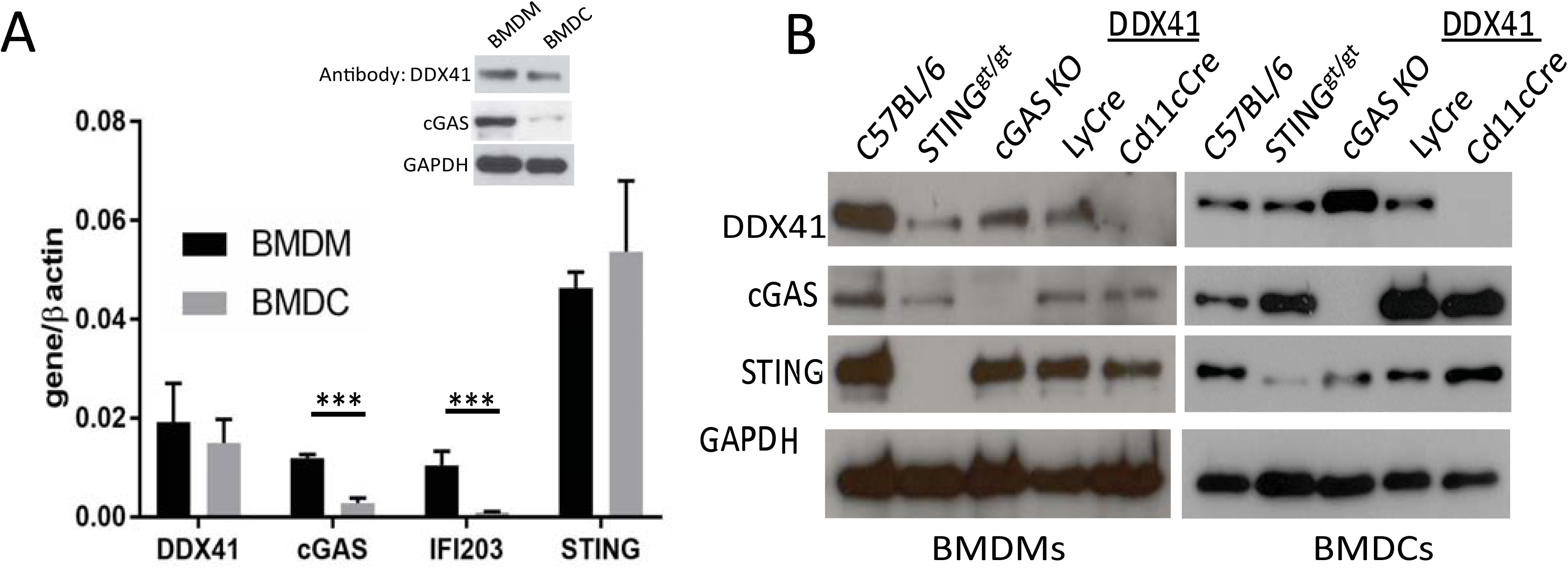
Characterization of DDX41 knockout BMMs and BMDCs. A) Basal expression of the different sensors in wild type BMDCs and BMDMs. Shown are the average and STDs of cells isolated from 3 different mice. Inset shows western blot of 40 μg each of extracts from BMDMs and BMDCs, probed with antisera against DDX41, cGAS and GAPDH. B) Relative expression of DDX41 in BMDMs and BMDCs. Forty μg of protein from cells isolated from mice of the indicated genotypes were analyzed by western blot with antisera to the indicated proteins. Gels are representative of 3 independent experiments. Averages from the 3 experiments are shown in in Supplementary Fig. 4B.

To determine whether DDX41 was important for IFN-induction in these cells types, we used mice with a knocked-in floxed DDX41 allele, in which the loxP sites flank exons 7 and 9 (Fig. S4A). We crossed these mice with CMV-Cre mice, but no complete KO pups were generated, suggesting that germline loss of Ddx41 causes embryonic lethality. We then crossed these mice with CD11cCre and LyCre transgenic mice to generate DC- and myeloid lineage-specific KOs, respectively. We also used BMDMs and BMDCs from cGas KO mice and Sting^gt/gt^ mice, which encode a mutant STING protein incapable of signaling and whose protein levels are greatly reduced (32). BMDMs from the LyCre-DDX41 and BMDCs from the CD11cCre-DDX41 mice were deficient in DDX41 RNA and protein but had wild type levels of STING and cGAS (Figs. 5B and S4B). Additionally, DDX41 protein levels were significantly higher in the cGas KO BMDCs but not BMDMs (Figs. 5B and S4B). Basal levels of IFN were not also not affected by loss of DDX41 (Fig. S4C) and FACS analysis demonstrated that DDX41 deficiency did not affect overall percentages of peripheral blood DCs or macrophages in the CD11cCre-DDX41 or LyCre-DDX41 mice (Fig. S4D). To ensure that DDX41 loss did not affect all innate immune responses, BMDCs and BMDMs from CD11cCre-DDX41 and LyCre-DDX41 mice were treated with the Toll-like receptor (TLR) 4 ligand LPS, the TLR3/MAVS pathway ligand poly (I:C) and cGAMP; cells from cGas KO, Sting^gt/gt^ and C57BL/6N mice served as controls. The response to LPS and poly (I:C) were similar to wild type in CD11cCre-DDX41 BMDCs, LyCre-DDX41 BMDMs, cGas KO and Sting^gt/gt^ BMDMs and BMDCs (Fig. S4E). cGAMP responses were reduced only in STING^gt/gt^ cells, as previously reported (33).

We then used these cells to examine the response to MLV infection. BMDCs or BMDMs lacking DDX41 mice showed little or no increase in type I IFN RNA (Fig. 6A) or protein (Fig. S5A) in response to MLV, even when TREX1 levels were reduced by siRNA treatment. BMDCs and BMDMs from Sting^gt/gt^ and cGas KO mice also had an abrogated antiviral IFNβ response under the same conditions (Figs. 6A, S5A). We also tested whether the response to HIV-1 was defective in the various mouse knockout cells, using pseudoviruses bearing the ecotropic MLV envelope; the IFNβ RNA response to HIV was diminished in both DDX41 and cGAS KO BMDMs and BMDCs (Fig. 6B). Thus, both sensors are required for the full type I IFN response to both MLV and HIV in mouse cells.

**Fig. 6.**
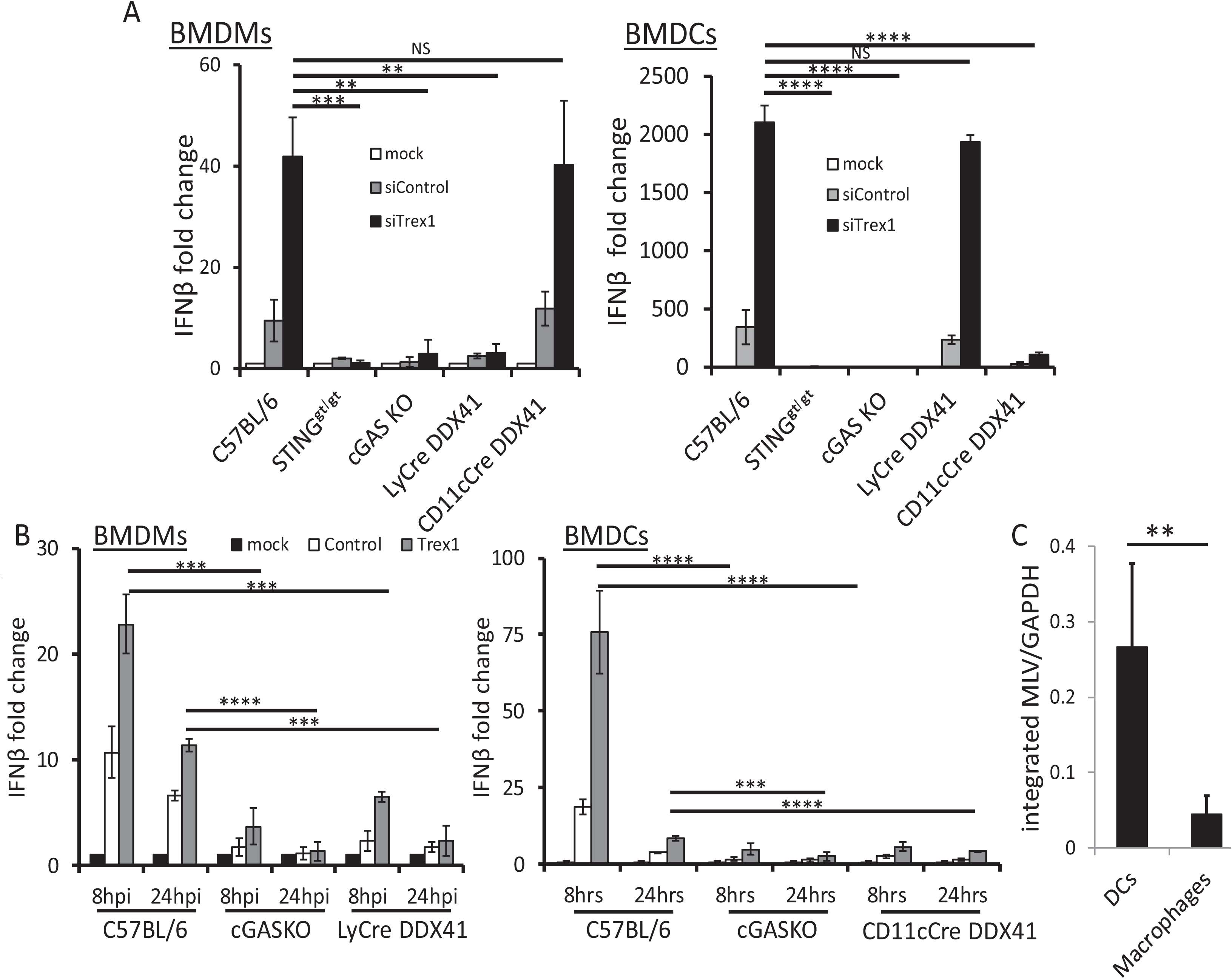
MLV and HIV induce a DDX41-dependent IFNβ response in BMDMs and BMDCs. A) BMDMs and BMDCs isolated from mice of the indicated genotypes were infected with MLV and at 2 hr pi, IFNβ levels were measured. The data in the graph are the average of 3 different experiments, each with macrophages and DCs from a different mouse. *, p≤ 0.05; **, p≤ 0.01; ***, p≤ 0.001 (unpaired T-test). Verification of the knockdown is shown in Fig. S5B. ELISAs measuring IFNβ protein are shown in Fig. S5A. B) HIV pseudotype infection in cGAS and DDX41 KO BMDMs and relative infection of BMDMs and BMDCs by MLV. HIV cores pseudotyped with ecotropic MLV glycoproteins were used to infect BMDMs (left panel) or BMDCs (right panel) from mice of the indicated genotypes. Trex1 knockdowns are shown in the right panels. Shown are the average of 3 independent experiments. ***, p≤ 0.001; ****, p≤ 0.0001 (unpaired T-test). C) DCs and macrophages were isolated from MLV-infected mice at 16 dpi and levels of integrated MLV were determined by qPCR. **, p≤ 0.01. Mock denotes mock-infected cells.

The TREX1-/DDX41-dependent IFNβ response to MLV infection was much higher in BMDCs than in BMDMs; there was a 2000-fold increase in IFNβ RNA in BMDCs compared to BMDMs, where the response was about 40-fold (compare y axes in Fig. 6A and 6B). To determine if this was due to increased infection, we isolated splenic DCs and macrophages from MLV-infected C57BL/6 mice at 16 days post-inoculation (dpi), as well as BMDCs and BMMs from mice of all the genotypes infected *ex vivo* with MLV. Integrated MLV DNA was analyzed by qPCR with a B1 repeat- and MLV LTR-specific primers. MLV infection of DCs was about one order of magnitude higher than that of macrophages, after either *in vivo* and *ex vivo* infection, independent of the mouse genotype (Figs. 6C and S6, respectively). Thus, while macrophages can be infected, sustain reverse transcription and mount a response to viral nucleic acids, DCs are more infected and respond more robustly to infection.

### Full suppression of MLV infection *in vivo* requires both DDX41 and cGAS

To determine whether DDX41 and cGAS functioned *in vivo* to suppress infection, we subcutaneously inoculated the CD11cCre-DDX41 and cGas KO mice with MLV and measured infection levels in the draining lymph node; wild type (DDX41f/f mice without Cre) and Sting^gt/gt^ mice served as controls. The CD11cCre-DDX41 and cGAS knockout mice showed significantly higher levels of infection than the wild type mice, while Sting^gt/gt^ mice had the highest level of infection (Fig. 7A).

Next, we tested whether DDX41 and cGAS acted synergistically *in vivo*. We treated CD11cCre-DDX41 and wild type mice with cGAS siRNAs and cGas knockout and wild type mice with DDX41 siRNAs; mice injected with the *in vivo* transfection reagent Invivofectamine alone served as controls. At 48 hr post-siRNA treatment, the mice were infected with MLV in the same footpad and at 24 hpi, RNA was isolated from the draining lymph node and examined for MLV RNA levels (Fig. 7A) and the extent of gene knockdown (Fig. S7). CD11cCre-DDX41 mice that received the cGAS siRNA and cGAS knockout mice that received the DDX41 siRNA were infected at >8-fold higher levels than wild type mice receiving no siRNA and at >3-fold higher levels than wild type mice receiving the DDX41 or cGAS siRNA. Infection levels in the CD11cCre-DDX41/cGAS siRNA group were not statistically different than the cGAS KO/DDX41 siRNA group (Fig. 7A). Wild type mice receiving the DDX41 or cGAS siRNAs were >2-fold more infected than untreated wild type mice and were not statistically different from each other. Sting^gt/gt^ mice had the highest levels of infection, about 2-fold higher than cGAS KO/DDX41 siRNA or CD11cCre-DDX41/cGAS siRNA mice.

**Fig. 7.**
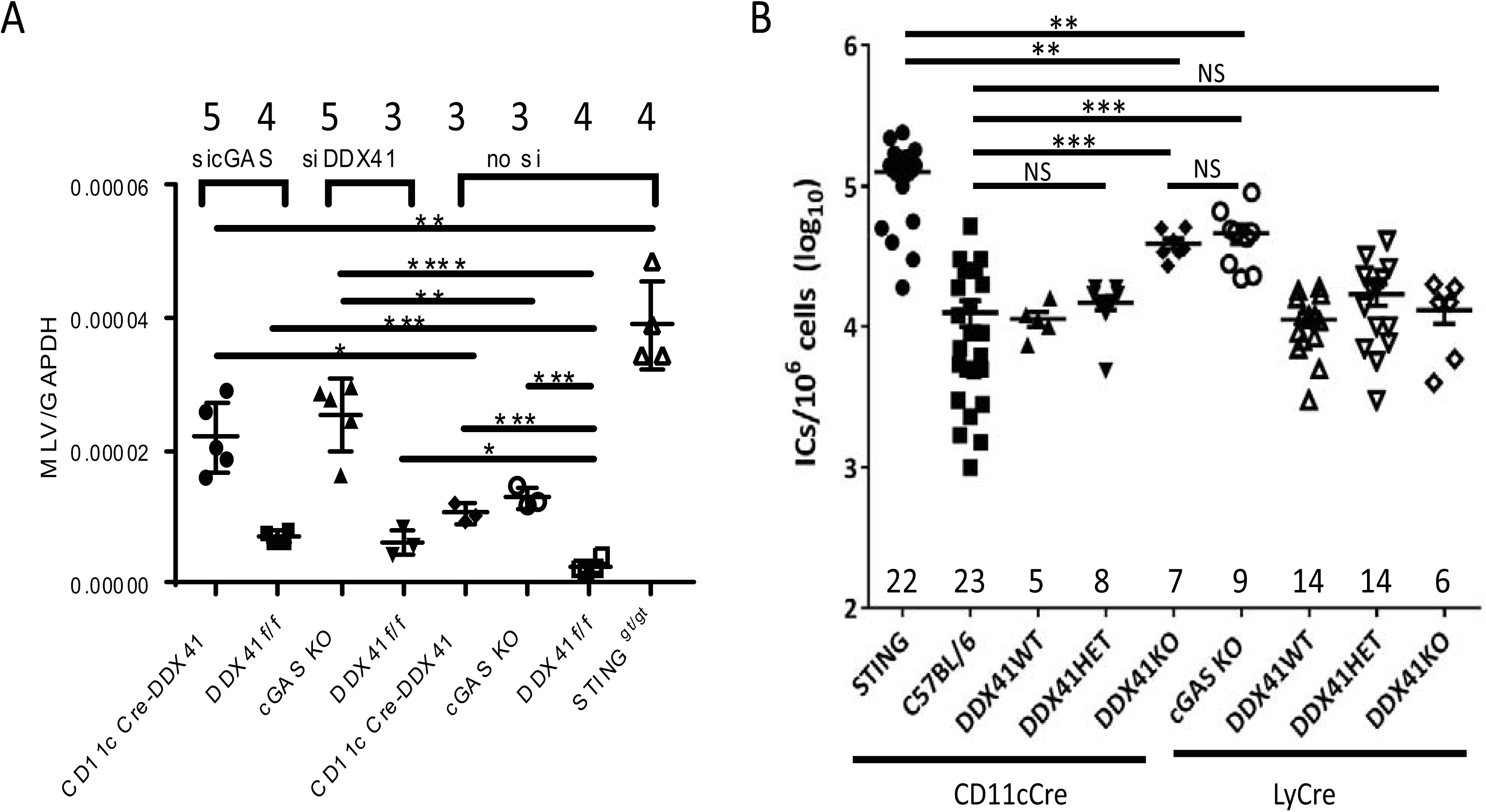
*In vivo* control of retrovirus infection requires both cGAS and DDX41. A) Mice of the indicated genotype received footpad injections of siRNAs. Forty-eight hrs post-injection, the mice were injected with of MLV. RNA was harvested from the draining lymph node at 24 hpi and analyzed for MLV. Knockdown verification is shown in Fig. S7. ****, p≤0.00001; ***, p≤0.0001; **, p≤0.001; *, p≤0.01B). (unpaired T-test). Number of mice in each group is shown above the graph. B) Newborn mice of the indicated genotypes were inoculated with MLV. At 16 days pi, virus titers in spleen were measured. Each point represents an individual mouse. The numbers of mice analyzed in each group is shown on the graph; each of the groups of mice came from 4 – 10 independent litters. Horizontal bars represent the average. **, p≤0.001, NS, not significant (Mann Whitney T-test).

We also examined whether DDX41 expression in BMDMs or BMDCs was important to suppress long term *in vivo* infection. Newborn offspring from crosses between LyCre-DDX41+/- and CD11cCre-DDX41 +/- mice as well as newborn C7BL/6, Sting^gt/gt^ and cGAS KO pups were inoculated with MLV and at 16 dpi, virus titers in their spleens were measured; this timepoint has been used extensively by us and others to examine MLV infection (4, 5, 7, 9, 34, 35). The genotyping of the intercrossed mice was carried out subsequent to measuring the virus titers. We thus compared infection levels between mice with total lack of DDX41 due to full knockout of the gene in the specific compartment to mice with only one knockout allele and to mice with no knockout of DDX41 (Fig. S6B).

Mice with complete knockout of DDX41 in DCs showed 5-fold higher infection than either wild type mice or mice heterozygous for the DDX41 knockout allele in this cell type (Fig. 6). cGas KO mice were also more infected, also to about 5-fold higher levels than wild type mice and the level of infection was the same as the CD11cCre-DDX41 mice (Fig. 6). In contrast, Sting^gt/gt^ mice were most highly infected with MLV, about 10-fold higher than wild type mice (Fig. 6). This confirms that cGAS and DDX41 are both required for full sensing of retroviral reverse transcripts and for the control of virus *in vivo*. Surprisingly, MLV infection of mice with complete knockout of DDX41 in macrophages was the same as wild type or heterozygotes (Fig. 6). Thus, although DDX41 sensed MLV infection in macrophages *in vitro* and *ex vivo*, this response *in vivo* was not sufficient to control infection.

## Discussion

The host factor APOBEC3, which both blocks reverse transcription and causes lethal mutation of the viral genome, is likely the first lines of defense against retroviruses, although incoming retroviruses do generate ligands that activate the innate immune system (9). We recently proposed that the major role for cytosolic sensing of reverse transcripts that escape the APOBEC3-mediated reverse transcription block is to induce expression of ISGs, including APOBEC3 itself (9). The most highly studied of these sensors, cGAS, is clearly a critical component of the foreign DNA recognition pathway, leading to STING activation and the type I IFN response. However, the role of other sensors implicated in the response to DNA generated during pathogen infection remains controversial. These include DDX41 as well as members of the ALR family (4, 10, 13, 36, 37). Here we show that DDX41 is a critical sensor of viral nucleic acids generated during reverse transcription and is required to control *in vivo* infection.

Retroviruses are unique in generating multiple different forms of nucleic acid during their replication in the cytoplasm which can be recognized as “foreign” by the host cell. DDX41 is likely recognizing the DNA/RNA hybrid generated in the first step of retrovirus replication and cGAS the dsDNA generated after strand translocation. While we showed that DDX41 and cGAS KO BMDMs or BMDCs showed diminished responses to transfected synthetic dsDNA or DNA/RNA molecules, DDX41 preferentially precipitated RNaseH-sensitive, DNA/RNA hybrid reverse transcripts generated during MLV infection and only depletion of DDX41 specifically reduced the IFN response to the RNaseH mutant virus, which generates more RNA/DNA hybrids than does wild type MLV. In contrast, cGAS precipitated DNaseI-sensitive reverse transcripts and cGAS-depletion did not completely abrogate the type I IFN response to the RNaseH mutant generated at the 1^st^ step of reverse transcription. Whether the presence of the tRNA primer bound to DNA/RNA hybrids or dsDNA plays a role in the recognition by DDX41 or cGAS, respectively, is currently not know.

*DDX41* belongs to a family of RNA helicases, with distinct DEAD/H box (Asp-Glu-Ala-Asp/His) domains, whose members have been implicated in translation, ribosome biogenesis, nuclear-cytoplasmic transport, organelle gene expression and pre-mRNA splicing (38-40). *DDX41* was recently identified as a tumor suppressor gene in familial and sporadic myelodysplastic syndrome/acute myeloid leukemia (MDS/AML), as well as other hematological malignancies (21, 39, 41). In MDS/AML, DDX41 is thought to interact with spliceosomal components and alter splicing, resulting in the inactivation of tumor suppressor genes or alterations in the balance of gene isoforms, although whether this occurs through protein-RNA, protein-DNA or protein-protein interactions is not known. Our data showing that DDX41 interacts with RNA/DNA hybrids are consistent with the known ability of DEAD box proteins’ recognition of RNA and suggest that DDX41 may have evolved an anti-viral cytoplasmic activity that takes advantage of its unique ability to interact with both RNA and DNA, as well as proteins. Another DEAD-box helicase, DDX3, was also recently implicated in the sensing of HIV RNA in DCs (42). However, DDX3 sensed abortive RNA transcribed from integrated proviruses, whereas DDX41 sensing occurred in the presence of the integrase inhibitor raltegravir, confirming that it works at a very early step of infection.

A previous study suggested that DDX41 might be the initial cytosolic sensor in BMDMs and that type I IFNs induced by DDX41 sensing lead to increased expression of cGAS, which is an ISG (43). However, at 2 hr pi, cGAS- and DDX41-deficient cells showed similar decreased IFNβ RNA levels after MLV infection, suggesting that both sensors are needed for the initial response. We showed previously that DCs get infected by MLV (5) and here we demonstrate that DDX41 in DCs but not macrophages was required for *in vivo* control of virus infection. The innate immune response initiated by DDX41 sensing of MLV in DCs may be due to higher levels of infection than in macrophages or because DCs are more effective at initiating antiviral responses. Whether the cGAS-dependent response is also required to control *in vivo* infection primarily in DCs is not known. Nevertheless, the results presented here contradict studies suggesting that DCs do not get infected but serve only as carriers that deliver intact retroviral virions to lymphocytes (44-46).

Previous work suggested that only cGAS is important for sensing retroviruses via the STING pathway (12, 47). These studies used VSV G protein-pseudotyped HIV or MLV cores. Both HIV and MLV naturally enter cells from a neutral compartment and it is possible that the use of VSV G, which directs entry to an acidic compartment, might affect the accessibility of different sensors to the reverse transcription complex. Additionally, these studies tested embryonic fibroblasts or BMDMs. However, as we demonstrated previously and our *ex vivo* and *in vivo* studies here demonstrate, DCs are likely the important targets of retroviruses (5, 48). Indeed, we also show here that endogenous cGAS expression in DCs, the relevant cell type for controlling MLV infection *in vivo*, is ^~^4-fold lower than that seen in macrophages, which could also account for the differences in our results with previous studies. Similar differences in cGAS expression occur in human macrophages and DCs (49).

Finally, earlier studies did not examine the effects of the different sensors on *in vivo* infection. We show here that effective *in vivo* control of MLV infection via the STING pathway requires both DDX41 and cGAS. However, as we and others have shown, the retrovirus capsid likely protects the reverse transcription complex from host sensors and other restriction factors, including APOBEC3 proteins (9, 50). This may explain why mice lacking DDX41 or cGAS show only 5-fold higher infection than wild type mice; even STING-deficient mice show only 10-fold higher infection (Fig. 7) (9). Our data are consistent with a requirement for both DDX41 and cGAS, the former perhaps in complex with IFI203, to achieve the full antiviral IFN response to retroviral reverse transcripts not protected by capsid or blocked by APOBEC3 proteins. Whether DDX41 requires interaction with IFI203 to achieve maximum effect *in vivo* will also be important to determine; however, *Ifi203* shares a high degree of identity in the noncoding as well as coding regions with several other genes in the *ALR* locus, making a gene-specific knockout difficult to achieve (51). How nucleic acid-bound DDX41 activates STING is also not yet understood, although the two molecules are known to directly bind each other (9, 13)

Our data suggest that there are multiple cytosolic sensors that recognize the different types of nucleic acids generated during retrovirus infection. Understanding the initial host response to infection by retroviruses is critical to our ability to determine how these viruses establish persistent infection as well the discovery of novel approaches to intervene in these infections.

### Experimental Procedures

#### Mice

Mice were bred at the University of Pennsylvania and the University of Illinois at Chicago. DDX41-flp mice (C57BL/6N) were constructed by TaconicArtemis GmbH and were derived by the University of Pennsylvania Transgenic & Chimeric Mouse Facility from *in vitro* fertilization of C57BL/6N embryos with sperm from a single male. LyCre (B6.129P2-Lyz2tm1(cre)Ifo/J) and Sting^gt/gt^ (C57BL/6J-Tmem173gt/J) mice were purchased from the Jackson Laboratory. CD11cCre mice (B6.Cg-Tg(Itgax-cre)1-1Reiz/J) were provided by Yongwon Choi and cGAS knockout mice by Michael Diamond and Skip Virgin (52). Apobec3 knockout mice were previously described (53). cGas/Apobec3 double knockout mice were generated by inter-crossing the two strains. All mice were housed according to the policies of the Institutional Animal Care and Use Committee of the University of Pennsylvania and of the Animal Care Committee of the University of Illinois at Chicago; all studies were performed in accordance with the recommendations in the Guide for the Care and Use of Laboratory Animals of the National Institutes of Health. The experiments performed with mice in this study were approved by the U. Pennsylvania IACUC (protocol #801594) and UIC ACC (protocol #15-222).

#### FACS analysis and sorting

Peripheral blood mononuclear cells were stained with anti-mouse F4/80-FITC (Biolegend) and anti-mouse CD11c-PE (BD Bioscience) antibodies. Cells were processed using a Beckman Coulter Cyan ADP. Results were analyzed using FlowJo software.

#### Virus

Moloney MLV and MLV^glycoGag^ mutant viruses were harvested from stably infected NIH3T3 fibroblasts, as previously described (54). All virus preparations were titered on NIH3T3 cells and analyzed by RT-qPCR for viral RNA levels as previously described (4). To generate the RNaseH mutant virus, the D524N mutation previously described by Blain and Goff (27) was introduced into the WT MLV infectious clone p63.2 (55) by site-directed mutagenesis using the Quickchange II XL site directed mutagenesis kit (Agilent Technologies) and the primers 5’-ACCTGGTACACGAATGGAAGCAGTCTCTTAC-3’/5’-GTAAGAGACTGCTTCCATTCGTGTACCAGGT-3’; the mutation was verified by sequencing. The p63.2 and p63.2^D524N^ plasmids were transfected in 293T cells using Lipofectamine 3000 (Invitrogen). The media of the transfected cells were harvested 48 hours post-transfection, centrifuged at 500g for 10 minutes at 4°C, filtered through a 0.45μm fiter and treated with DNaseI recombinant RNase Free (Roche). Virus levels were determined by the QuickTiter^™^ MuLV Core Antigen Elisa Kit (MuLV p30) (Cell Biolabs, Inc) and by titering on NIH 3T3 cells stably transfected with pRMBNB, which expresses the MLV *gag* and *pol* genes (28).

#### BMDM and BMDC cultures

BMDMs and BMDCs were isolated from hind limbs of 10- to 12-week-old cGas KO, Sting^gt/gt^, LyCre-DDX41, CD11cCre-DDX41 and C57BL/6 mice as previously described (56). BMMs were cultured in DMEM supplemented with 10% FBS, 10ng/ml Macrophage Colony Stimulating Factor (Invitrogen), 1 mM sodium pyruvate, 100 U/ml penicillin and 100 μg/ml streptomycin and were harvested 7days after plating and were seeded in 96-well plates for infection assays. BMDCs were cultured in RPMI supplemented with 5% FBS and differentiated with recombinant murine granulocyte-macrophage colony-stimulating factor (20 ng/ml; Invitrogen). Both procedures result in cultures that are ^~^80% - 85% pure.

#### cGAMP stimulation of macrophages

Knockdowns with the indicated siRNAs were performed in NR9456 macrophages (immortalized macrophage cell line derived from C57BL/6 wild type mice) (57) (BEI Resources, NIAID, NIH) using RNAiMAX, as previously described (9). The next day, cells were transfected with Lipofectamine 2000 (Invitrogen) and 4ug of cGAMP (Invivogen), 16 hrs later the cells were infected with MLV^glycoGag^ and harvested 2 hpi. RNA isolation and qPCR analysis was performed as previously described (9).

#### Virus infection of macrophages and DCs

NR9456 macrophages, BMDMs and BMDCs were siRNA-transfected. Forty-eight hrs after transfection, the cells were infected with wild type, MLV^glycoGag^ or D542N mutant M-MLV (MOI of 2) and harvested at the indicated times after infection. For some experiments, the cells were treated with 200nM raltegravir for 2 hr prior to infection and then infected with MLV^glycoGag^ virus in the presence of drug. Cells were harvested 2 hpi; RNA isolation and RT-PCR were performed as previously described (9). Primers used for detection of actin, Trex1 and IFNβ were previously described (4, 58). Primers to amplify the MLV 2LTR closed circles are 5’-GAGTGAGGGGTTGTGGGCTCT-3’/5’-ATCCGACTTGTGGTCTCGCTG-3’ (59). Primers used to amplify late reverse transcripts (P_3’R_/P_3’L_) are 5’-TAACGCCATTTTGCAAGGCA-3’/5’-GAGGGGTTGTGGGCTCTTTT-3’; strong-stop DNA primers were reported previously (4).

#### BMDM and BMDC treatment with synthetic ligands

BMDMs and BMDCs isolated from C57BL/6, Sting^gt/gt^, cGas KO, LyCre-DDX41, CD11c DDX41 mice were transfected with 2 ng/μl poly IC (Sigma) and 4 ng of cGAMP using Lipofectamine 3000; cells were also treated with 1 ng/μl LPS (Sigma). The cells were harvested at 6 hr post-treatment. RNA was isolated and cDNA was generated using Superscript III kit (Invitrogen). RT-PCR was performed to measure IFNβ RNA levels, as previously described (4).

#### siRNA knockdown and knockdown verification

NR-9456, BMDMs and BMDCs were transfected with the indicated siRNAs (9) using Lipofectamine RNAiMAX reagent (Invitrogen). RNA was isolated using the RNeasy Mini Kit (Qiagen). All siRNAs used in this study were previously shown to be on-target and to decrease both RNA and protein levels (9). cDNA was made using the SuperScript III First Strand Synthesis System for RT-PCR (Invitrogen). RT-PCR was performed using the Power SYBR Green PCR master mix kit (Applied Biosystems). Primers for the verification of the knockdowns have been previously described (4, 9).

#### IFNβ ELISAs

BMDMs and BMDCs were transfected with a control- or a Trex1-specific siRNA using Lipofectamine RNAiMAX reagent (Invitrogen). Cells were then infected with MLV^glycoGag^ and the culture media was harvested 4 hpi. The levels of IFNβ in the culture media were measured using the LEGEND MAX^TM^ Mouse IFNβ ELISA kit (Biolegend) per manufacturer’s recommendation.

#### HIV pseudoviruses

Retroviral vectors bearing the Moloney MLV Env and HIV (pNL4-3) cores were produced by transient transfection into 293T cells using Lipofectamine 3000 (Invitrogen), as previously described (9). Pseudoviruses were harvested at 48 hpi and the pseudoviruses were treated with DNaseI (20u/ml for 45min at 37°C) (Roche) and concentrated using AMICON columns.

#### Nucleic acid pulldowns

DDX41myc/his, IFI203-HA and cGAS-V5 plasmids have been previously described (9, 60). 293MCAT cells transfected with pcDNA3.1 (empty vector), cGAS-V5, and DDX41myc/his, were infected with virus and at 4 hpi, the cells were cross-linked with 1% formaldehyde in media. Cross-linking was quenched with 2.5M glycine, extracts prepared and then incubated overnight with anti-myc-or anti-HA-agarose beads (Sigma) or anti-V5 antibody (Invitrogen) with G/A-agarose beads (SantaCruz). The beads were washed with high-salt buffer (25mM Tris-HCl, pH 7.8, 500mM NaCl, 1mM EDTA, 0.1% SDS, 1% TritonX-100, 10% glycerol) and with LiCl buffer (25mM Tris-HCl, pH 7.8, 250mM LiCl, 0.5% NP-40, 0.5% Na-deoxycholate, 1mM EDTA, 10% glycerol). The immunoprecipitated nucleic acid was eluted from the beads at 37°C in 100mM Tris-HCl, pH 7.8, 10mM EDTA, 1% SDS for 15 min and the protein-nucleic acid cross-linking was reversed by overnight incubation at 65°C with 5M NaCl. The eluted nucleic acid was purified using the DNeasy Kit (Qiagen) and analyzed with RT-PCR strong stop primers (primers P_R_ and P_U5_ in Fig. 3A) or 3’LTR primers P_3’R_-P_3’L_ in Fig. 4A) (4). For analysis of the tRNA-bound MLV nucleic acid, the same procedure was used, except the eluted nucleic acid was reverse transcribed prior to PCR with the P_R_ primer and another primer that annealed to nt 39-57 in tRNA^pro^ (P_tRNA_ in Fig. 3A) (5’-GCTCTCCAGGGCCCAAGTT-3’)(61). For the nuclease treatments, after the nucleic acids were released from the protein cross-link, they were ethanol precipitated and treated at 37°C with 50u RNase A (Thermo) for 20min in the presence of 300mM NaCl, 4u DNase I (Roche) with the reaction buffer provided with the enzyme for 20min or 3u of RNase H (Thermo) for 20min in the reaction buffer provided with the enzyme. Samples were digested with proteinase K and phenol-chloroform extracted and the nucleic acids were subjected to qPCR analysis as described earlier in this section.

#### MLV infection levels

NR9456 cells were infected with WT or D542N virus and 2hrs hpi, cellular DNA and RNA were isolated. RNA was reverse transcribed using a Superscript III kit (Invitrogen) and the resultant cDNA was used for quantitative PCR using the P_R_-P_tRNA_ primers. DNA was subjected to quantitative PCR using the P_R_-P_U5_ and the P_3’LTRF_-P_3’LTRR_ primers. Bone marrow from C57BL/6 mice was isolated and differentiated to BMDMs and BMDCs. BMDCs and BMDMs were infected with MLV (0.1 MOI/cell). Cells were harvested at 24 and 48 hpi. For analysis of cell subset infection in vivo, newborn mice were infected i.p. with MLV. At 16 dpi, splenocytes were isolated and FAC-sorted directly into 15ml collection tubes using a MoFlo Astrios^™^ (Beckman Coulter, Brea, CA) at the UIC Cell Sorting Facility; anti-F4/80-FITC and -CD11c-PE was used to distinguish macrophages and DCs, respectively. DNA was isolated by using the DNeasy Kit (Qiagen). Quantitative PCR was performed to measure integrated MLV DNA using the following primers: 5’-CCTACTGAACATCACTTGGGG-3’/5’-GTTCTCTAGAAACTGCTGAGGGC-3’ and normalized to GAPDH.

#### Western Blots

Protein extracts from the BMDMs and BMDCs were run on 10% SDS-polyacrylamide gels and transferred to PVDF Immobulon membranes (Thermo). Rabbit anti-STING, anti-cGAS, anti-phosphoIRF3 (Ser 396), anti-IRF3, anti-TBK1 and anti-phosphoTBK1 (Ser 172) and HRP-conjugated anti-rabbit antibodies, all from Cell Signaling Technology, mouse monoclonal anti-DDX41 (SantaCruz Biotechnology) and -mouse antibody (Sigma Aldrich) were used for detection, using either ECL Western blot detection reagent or ECL prime Western blot detection reagent (GE Healthcare Life Sciences).

#### *In vivo* siRNA knockdown

siRNAs were purchased from Ambion (Life Technologies). The Invivofectamine 3.0 Starter Kit (Invitrogen, Life Technologies) was used according to the manufacturer’s protocol. Each siRNA solution (2.5nmol/μl) was combined with complexation buffer and Invivofectamine reagent for 30 minutes at 50°C. Footpad injections of the siRNA/Invivofectamine complex or Invivofectamine alone were carried out 48h prior to infection with MLV (2.5 × 10^5^ IC/mouse) in the same footpad. Each mouse received 20nmol of siRNA. After 24 hpi, mice were euthanized and draining lymph node tissues were collected and harvested for RNA isolation. MLV RNA levels were measured by RT-qPCR, as previously described (34). Knockdown of the siRNA-targeted gene was also verified by RT-qPCR as described above.

#### *In vivo* infections

For systemic infections, two-day old mice (C57BL/6N, cGAS KO, STING^gt/gt^ and the tissue-specific DDX41 KO mice described in Suppl. Table 1) were infected intraperitoneally with 2×10^4^ infectious center (IC) units of MLV and then harvested at 18 days pi and virus titers in spleens were measured by IC assays, as previously described (4). The *in vivo* infection studies were performed both at the University of Pennsylvania and the University of Illinois. The DDX41 knockout mice were housed side-by-side with the STING^gt/gt^ and cGAS mice, and crossed with BL/6N mice from our colony.

#### Statistical Analysis

Each experiment was done with 3 technical replicates/experiment. Data shown is the average of at least 3 independent experiments, or as indicated in the Fig. legend. Statistical analysis for the various experiments was performed using the GraphPad/PRIZM software. All raw data are deposited in Mendeley dataset accession http://dx.doi.org/10.17632/j4mgm4v9t3.1.

## Acknowledgements

We thank Gerard Zurawski for help in obtaining the DDX41-flox mice, Skip Virgin and Mike Diamond for the cGAS KO mice, Yongwon Choi for the CD11cCre mice, the Transgenic and Chimeric Mouse Facility of the University of Pennsylvania for generating the DDX41-flox mice by *in vitro* fertilization, Lorraine Albritton for the Moloney MLV construct, Vinay Pathak for the gag-pol (pRMBNB) construct and David Ryan for help with maintaining the different mouse lines. Raltegravir was obtained through the NIH AIDS Reagent Program and NR9456 cells were obtained from the BEI Resources, NIAID, NIH.

## Supporting Information

**Fig. S1**. Knockdown verification. Related to Fig. 1. A) Knockdown of genes in Fig. 1A. The knockdown of the three target genes (DDX41, IFI203 an STING) were all done in the presence of Trex1 knockdown, as indicated in the Fig. 1 legend. B) Knockdown of genes in Fig. 1C. See Fig. 1 legend for details.

**Fig. S2**. DDX41 senses DNA in the cytoplasm via its DEAD domain. Related to Fig. 2. A) Increasing nuclear dsDNA by inhibiting proviral DNA integration has no effect on DDX41-mediated sensing. NR9456 cells were pretreated with raltegravir (200nM) and then infected with MLV (MOI=2) in the presence of drug. DNA and RNA were isolated from infected cells 2 hpi and analyzed for unintegrated viral DNA (2LTR) or IFNβ RNA levels. Values are shown as mean ± STDs of three experiments. P values were determined by an unpaired T-test. (*, p≤ 0.05; **, p≤ 0.01; ***, p≤ 0.001). Inset shows levels of Trex1 RNA knockdown.

**Fig. S3**. Knockdown verification. Related to Fig. 4. A) Knockdown of genes in Fig. 4C. See Fig. 4 legend for details.

**Fig. S4**. Characterization of DDX41 KO mice. Related to Fig. 5 and 6. A) Map of the DDX41 locus and inserted lox P sites. Expression of Cre recombinase results in the deletion of exons 7 - 9. B) Quantification of DDX41, cGAS and STING protein in various knockout mouse cells. Shown is the mean +/− STD for 3 independent western blots of cells from 3 different mice of each strain. *, p ≤ 0.05 compared to BL/6, STING^gt/gt^ and CD11cCreDDX41 (unpaired T-Test). C) Basal IFNβ RNA levels in DDX41 KO BMDMs and BMDCs. RNA was isolated from BMDMS of 3 mice and BMDCs of 2 mice each of the indicated genotypes and qPCR was performed for IFNβ levels, using a standard curve to measure relative levels. Shown is the mean +/− STD. Abbreviations: ND, not done. D) PBMCs from 4 mice of each genotype were stained with conjugated anti-CD11c (DCs) or anti- F4/80 (macrophages) antibodies and analyzed by FACS. E) Treatment of DDX41 KO BMDMs and BMDCs with different ligands. BMDMs and BMDCs from the cGAS, LyCre DDX41 and CD11cCre-DDX41 KO and STING^gt/gt^ mice were treated with the indicated ligands, as described in Supplemental Experimental Procedures. RNA was isolated after 6 hr of treatment for all ligands and subjected to RT-qPCR. Shown is the average of 2 experiments (triplicate technical replicates) with cells isolated from different mice.

**Fig. S5**. Loss of DDX41 decreases the IFN response to MLV and HIV infection. Related to Fig. 6. A) BMDMs and BMDCs from C57, cGAS, LyCre-DDX41 and CD11cCre-DDX41 were treated siCont or siTrex and then infected with MLV^glycoGag^. Four hrs later supernatants were used to perform an ELISA for IFNβ levels. Shown are the average of 3 independent experiments. ****, p≤ 0.0001 (unpaired T-test). B) Verification of Trex1 knockdown in BMDCs and BMDMs in Fig. 6A. C) Verification of the knockdowns in BMDMs and BMDCs in Fig. 6B.

**Fig. S6**. BMDCs are more infected by MLV than BMMS. Related to Fig. 6. BMDMs and BMDCs from mice of the indicated genotypes were infected with MLV and at 24 and 48 hr pi infection, DNA was isolated from cells and subjected to qPCR, using one primer to mouse genomic DNA and the other to the viral long terminal repeat. Values are shown and mean ± SEM of 2 experiments done for each cell type. highlighted boxes refer to homozygous and white boxes to heterozygous DDX41 tissue-specific knockouts, gray boxes refer to WT mice.

**Fig. S7**. *In vivo* control of retrovirus infection requires both cGAS and DDX41. Related to Fig. 7. A) Knockdown verification of the in vivo siRNA experiment presented in Fig. 7A. B) Genotypes of the mice tested for MLV infection in Fig. 7B. Parental mice were Cre+/−, DX41+/−. Cre refers to either LyCre or CD11cCre, as indicated in the text. Yellow highlighted boxes refer to homozygous and white boxes to heterozygous DDX41 tissue-specific knockouts, gray boxes refer to WT mice.

